# Blind prediction of noncanonical RNA structure at atomic accuracy

**DOI:** 10.1101/223305

**Authors:** Andrew Watkins, Caleb Geniesse, Wipapat Kladwang, Paul Zakrevsky, Luc Jaeger, Rhiju Das

## Abstract

Prediction of RNA structure from nucleotide sequence remains an unsolved grand challenge of biochemistry and requires distinct concepts from protein structure prediction. Despite extensive algorithmic development in recent years, modeling of noncanonical base pairs of new RNA structural motifs has not been achieved in blind challenges. We report herein a stepwise Monte Carlo (SWM) method with a unique add-and-delete move set that enables predictions of noncanonical base pairs of complex RNA structures. A benchmark of 82 diverse motifs establishes the method’s general ability to recover noncanonical pairs *ab initio*, including multistrand motifs that have been refractory to prior approaches. In a blind challenge, SWM models predicted nucleotide-resolution chemical mapping and compensatory mutagenesis experiments for three *in vitro* selected tetraloop/receptors with previously unsolved structures (C7.2, C7.10, and R1). As a final test, SWM blindly and correctly predicted all noncanonical pairs of a Zika virus double pseudoknot during a recent community-wide RNA-puzzle. Stepwise structure formation, as encoded in the SWM method, enables modeling of noncanonical RNA structure in a variety of previously intractable problems.

## Introduction

Significant success in protein modeling has been achieved by assuming that the native conformations of a macromolecule have the lowest free energy and that the free energy function can be approximated by a sum of hydrogen bonding, van der Waals, electrostatic, and solvation terms that extend over Angstrom-scale distances. Computational methods that subject large pools of low-resolution protein models to all-atom Monte Carlo minimization guided by these free energy functions have achieved near-atomic-accuracy predictions in the CASP community-wide blind trials (*1*). When adapted to RNA structure modeling, analogous methods have consistently achieved nucleotide resolution in the RNA-Puzzle blind trials but have not yet reached atomic accuracy, aside from previously solved motifs that happen to recur in new targets (*2*). A disappointing theme in recent RNA-Puzzle assessments is that the rate of accurate prediction of noncanonical base pairs is typically 20% or lower, even for models with correct global folds (*2*). Without recovery of such noncanonical pairs, RNA computational modeling will not be able to explain evolutionary data, predict molecular partners, or be prospectively tested by compensatory mutagenesis for the myriad biological RNAs that are being discovered at an accelerating pace.

The lag between the protein and RNA modeling fields is partly explained by differences in how protein and RNA molecules fold. Protein structures are largely defined by how αhelices and β-sheets pack together. As abundant data exist on these regular protein elements and their side-chain interactions, protein models with reasonable accuracy can often be assembled from fragments of previously solved structures. Less regular loops interconnecting α- and β-elements are less critical for defining protein folds. Indeed, those loops are typically not recovered at high accuracy, even in the most exceptional blind predictions (*3-5*). In contrast, the predictable and geometrically regular elements of RNA folding are Watson-Crick helices that sequester their side chains and therefore cannot be positioned by direct side-chain interactions. Instead, the RNA loops interconnecting those helices form intricate noncanonical base pairs that define an RNA’s global helix arrangement. The RNA structure prediction problem, more so than the protein problem, depends on discovering these irregular loop conformations and their associated noncanonical base pairs *ab initio*. Unfortunately, discovering the lowest free energy conformations of new noncanonical loop motifs has not generally been tractable due to the vast number of deep, local minima in the all-atom folding free energy landscape of even the smallest such motifs. Essentially all 3D RNA modeling methods, including MC-Sym/MC-Fold, Rosetta FARFAR, iFoldRNA, SimRNA, and VFold3D, use coarse-grained modeling stages that allow for smoother conformational search but generally return conformations too inaccurate to be refined to atomic accuracy by Monte Carlo minimization or molecular dynamics refinement (*6-10*).

To address this challenge, we have developed Rosetta methods that attempt to remove barriers in conformational search through addition of residues one-at-a-time rather than through low resolution coarse-graining or through small perturbations to fully built conformations. We previously described how step-by-step buildup of an RNA structure, enforcing low-energy conformations for each added nucleotide, could lead to atomic accuracy models of irregular single-stranded RNA loops (*11*). The calculation, instantiated in the Rosetta modeling framework, involved a deterministic enumeration over buildup paths, analogous to classic dynamic programming methods developed for canonical RNA secondary structure prediction (*11, 12*). This enumerative stepwise assembly (SWA) method guaranteed a unique solution for the final conformational ensemble but necessitated large expenditures of computational power. For example, calculations for even small loops of 5-7 nucleotides required tens of thousands of CPU hours (*11*); junctions involving multiple interacting strands would further increase computational cost to many millions of CPU hours, which is currently prohibitive.

In the hopes of reducing this computational expense, we hypothesized that the stepwise addition moves developed for SWA might still be effective at producing high accuracy models if implemented as part of a stochastic sampling scheme rather than deterministic enumeration. To test this hypothesis, we have developed stepwise Monte Carlo (SWM), a Monte Carlo optimization method whose primary moves are the stepwise addition moves of SWA. Here we report that SWM enables significant increases in the computational speed of *ab initio* structure prediction and describe applications of SWM to previously intractable noncanonical RNA structures. The tests include stringent blind evaluation through prospective experimental tests and the RNA-puzzle community-wide structure prediction challenge.

## Results

### Efficient implementation of stepwise Monte Carlo (SWM)

Fig. 1 illustrates the stepwise Monte Carlo procedure, which has been implemented in the Rosetta framework (*13*) and is also freely available through an online ROSIE server (Materials and Methods) (*14*). In realistic 3D RNA modeling problems, RNA helices are typically known *a priori* from secondary structure prediction methods. The main goal is therefore to infer lowest free energy conformations of loops that connect these helices, such as the four nucleotides GCAA closing a hairpin (Fig. 1A) or two strands, each with a single guanosine, in the GG mismatch motif (Fig. 1B). Our prior work (*11, 15, 16*) introduced stepwise addition moves that allow building of such nucleotides one at a time, starting from conformations with helices only (*Start*, Figs 1A-B). Conceptually, each addition was proposed to simulate the stepwise formation of well-defined structure from “random coil”-like ensembles (dotted lines, Fig. 1A-B).

**Figure 1.**
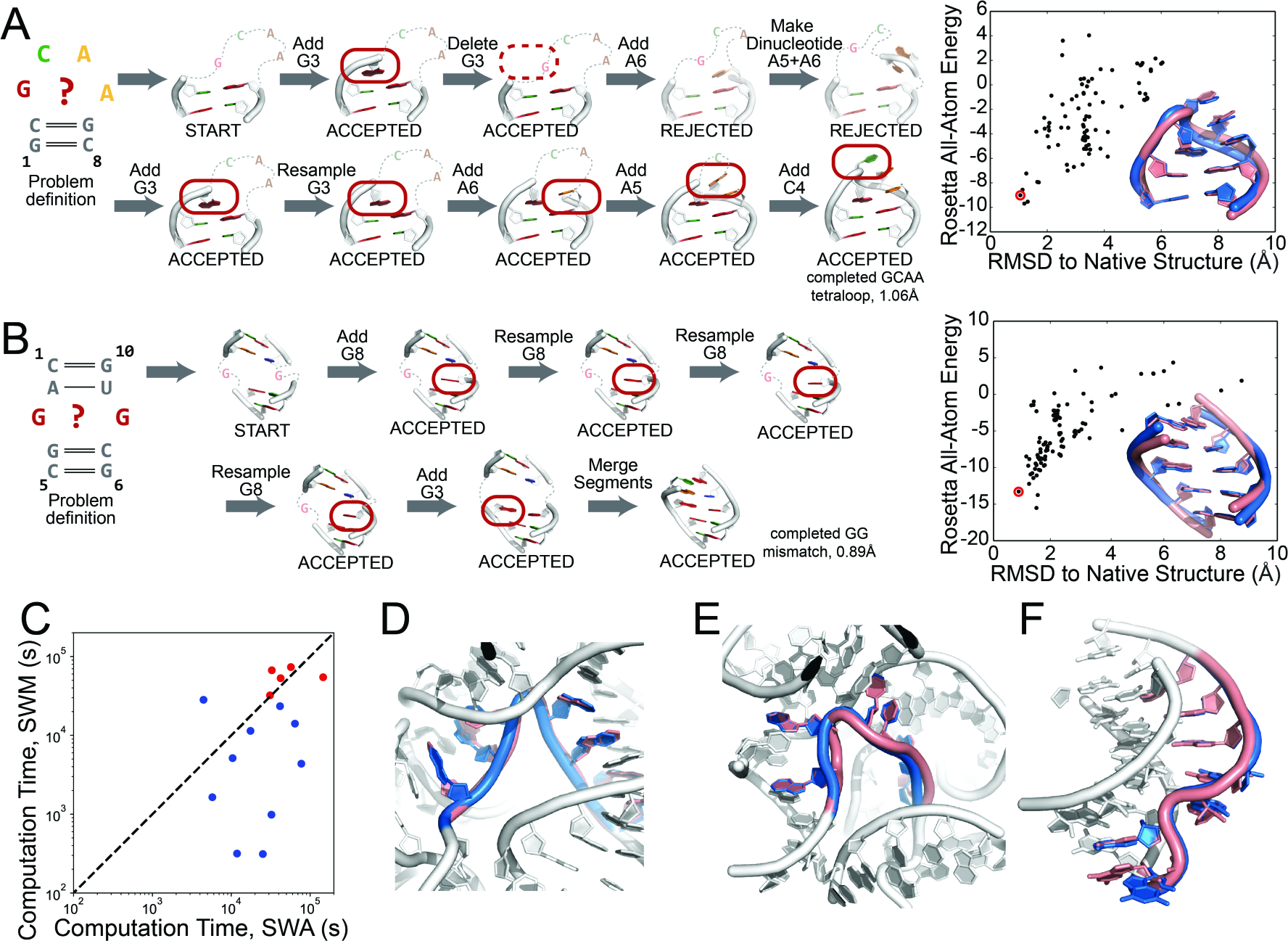
Stepwise Monte Carlo (SWM) efficiently searches the complex energy landscapes of noncanonical RNA loops. SWM trajectories solve **(A)** a GCAA tetraloop (PDB: 1ZIH) and **(B)** a two-strand GG-mismatch two-way junction (1F5G) in 10 moves or less (left panels). Final structures achieve low free energies and sub-Angstrom RMSD accuracies; numerous such structures appear in simulations involving 100 models (right-hand panels). **(C)** Significantly reduced CPU time is required for convergence of SWM compared to enumeration by SWA (*11*), except for loops drawn from the 23S rRNA (red). SWM models for **(D)** J2/3 from group II intron (3G78), modeled with prior energy function used for SWA, and **(E)** 23S ribosomal RNA loop (1S72) and **(F)** L2 loop of viral pseudoknot (1L2X), both modeled with updated Rosetta free energy function, illustrate sub-Angstrom recovery of irregular single-stranded loops excised from crystal structures.

Here, we carry out such addition moves stochastically, choosing at random positions on which to prepend or append new nucleotides (*Add*, Figs. 1A-B) rather than enumerating such additions at all possible positions (as was implemented previously (*11, 15*)). These stochastic moves are accepted if they lower the computed free energy of the model or if the free energy increment is lower than a thermal fluctuation energy as set by the Metropolis criterion (*17*). To maintain detailed balance in the Monte Carlo scheme, the moves intermix these additions with deletions of single residues, again chosen randomly and accepted based on the Metropolis criterion (*Delete*, Fig. 1A). These deletion moves simulate the transient unstructuring of nucleotides at the edges of loops. To allow buildup of multistrand motifs, we developed moves to merge and split independent regions of RNA, such as regions associated with the different helices of multi-helix junctions (*Merge*, Fig. 1B). Last, we allowed resampling of randomly chosen internal degrees of freedom, maintaining chain closure with robotics-inspired kinematic closure algorithms (*18, 19*) (*Resample*, Fig. 1A). These resampling moves could not be incorporated into the previous enumerative SWA due to the large number of increased modeling pathways that would need to be enumerated.

Before testing SWM, it was unclear whether a stochastic search might allow for efficient *ab initio* recovery of RNA loop conformations. Indeed, in our prior work on enumerative stepwise assembly we posited that “An inability to guarantee exhaustive conformational sampling has precluded the consistent prediction of biomolecular structure at high resolution” (*11*). Nevertheless, in our test cases, SWM did achieve efficient search over the free energy landscape, despite the lack of a guarantee of exhaustive conformational sampling. Figs. 1A-B illustrate the recovery of the sub-Angstrom accuracy conformations of the GCAA tetraloop and GG mismatch motifs, using less than three hours of computation on a single modern laptop computer to create 100 models. Furthermore, these runs were ‘convergent’: different simulations independently achieved the same low-energy configurations repeatedly (numerous models within 2 Å of experimental structures, right panels in Figs. 1A-B), suggesting highly efficient sampling. For comparison, modeling of these small loops by SWA enumeration required use of many thousands of CPU-hours due to the requirement of enumerating multiple loop conformations over multiple buildup pathways.

### *Ab initio* recovery of complex single-stranded RNA loops

After preliminary tests on simple loops, we carried out SWM modeling on a set of 15 single-stranded RNA loops excised from crystal structures, previously used to benchmark the enumerative SWA method (*11*). These loops were specifically chosen because of their irregularity; they each harbor non-A-form backbone conformations, form noncanonical pairs with surrounding residues, and span different helices in functional RNAs. We confirmed that for nearly all these trans-helix cases, SWM produces conformations that give computed free energies and accuracies as low as those achieved by SWA (median energy gap and RMSD values, Table S1; Fig. 1C-1D). In many cases, the computational cost for achieving convergent and accurate modeling was reduced by up to two orders of magnitude (Supplemental Methods, Fig. 1C). Exceptions to this speed increase were loops that needed to be rebuilt into the 23S ribosomal RNA (red points, Fig. 1C), which featured a particularly large number of surrounding nucleotides, and concomitantly many viable interacting conformations. This observation suggested that SWM would be particularly efficient for *ab initio* modeling of motifs that primarily form noncanonical interactions within the motif, as is typically the case for RNA junctions and tertiary contacts (see below), but not the longest ribosomal loops. Furthermore, through its increased speed, SWM allowed us to confirm that recent updates to the Rosetta free energy function (*20*) and estimation of conformational entropy of unstructured segments generally improved modeling accuracy for single-stranded RNA loops (Tables S1 and S2). The improvements included rescue of some of the ribosomal RNA loops (Fig. 1E) and solution of a loop from a beet western yellow virus frameshifting pseudoknot that was previously not solvable by SWA (Fig. 1F; Table S3) (*11*). Supplemental Text provides a more detailed description of energy function updates and results on this trans-helix loop benchmark.

### *Ab initio* recovery across complex noncanonical motifs

To evaluate SWM more broadly, we expanded the 15 single-stranded loop benchmark to a larger set of 82 complex, multi-stranded RNA motifs that we encountered in previous RNA-puzzles and other modeling challenges (Table S3 and Fig. S1). Due to the efficiency of SWM modeling, we could test a benchmark that was nearly three times larger than our most extensive previous efforts (*8*). Over the entire benchmark, SWM achieved a mean and median RMSD accuracy (over the top five cluster centers) of 2.15 and 1.49 Å and mean and median recovery of non-Watson-Crick pairs of 76% and 96%, respectively (Table 1, Table S3, and Fig. S2). We observed numerous cases in which the SWM model and experimental structure were nearly indistinguishable by eye (Fig. 2). Examples included two-stranded motifs that required orders of magnitude higher computational expense with the prior enumerative stepwise assembly method (*16*), such as the most conserved domain of the signal recognition particle (Fig. 2A; 1.26 Å RMSD, 5 of 5 noncanonical pairs recovered) and the first RNA-Puzzle challenge, a human thymidylate synthetase mRNA segment (Fig. 2B; 0.96 Å, 1 noncanonical pair and 1 extrahelical bulge recovered) (*2*). For several test cases, there was experimental evidence that formation of stereotyped atomic structures required flanking helices to be positioned by the broader tertiary context. If the immediately flanking helix context was provided, the median RMSD accuracy and non-Watson-Crick base pair recovery in these cases was excellent (1.19 Å and 100%; Table 1, Fig. S2 and Table S3), as illustrated by the J5/5a hinge from the P4-P6 domain of the *Tetrahymena* group I intron (*21*) (Fig. 2C; 0.55 Å RMSD, all 4 noncanonical pairs and all 3 extrahelical bulges recovered).

**Figure 2.**
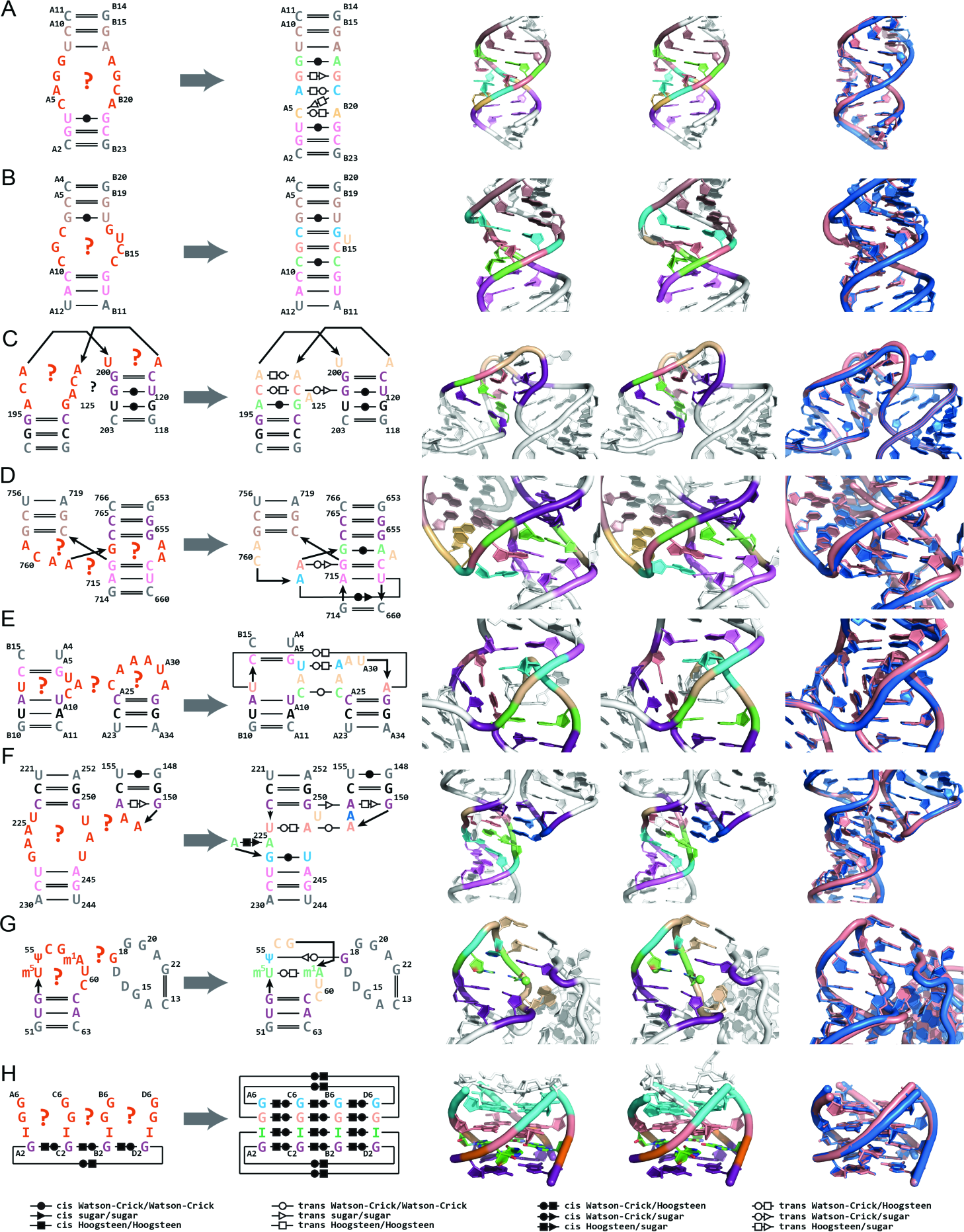
SWM recovers noncanonical base pairs *ab initio* for complex RNA motifs. From left to right in each panel, 2D diagram with problem definition, 2D diagram with experimental noncanonical base pairs, experimental 3D model, SWM 3D model, and 3D overlay (experimental, marine; SWM model, salmon). Motifs are **(A)** most conserved domain of human signal-recognition particle (PDB ID: 1LNT); **(B)** noncanonical junction from human thymidylate synthase regulatory motif, RNA-puzzle 1 (3MEI); **(C)** irregular J5/5a hinge from the P4-P6 domain of the *Tetrahymena* group I self-splicing intron (2R8S); **(D)** P2-P3-P6 three-way A-minor junction from the Varkud Satellite nucleolytic ribozyme, RNA-puzzle 7 (4R4V); **(E)** tertiary contact stabilizing the *Schistosoma* hammerhead nucleolytic ribozyme (2OEU); **(F)** tetraloop/receptor tertiary contact from the P4-P6 domain of the *Tetrahymena* group I self-splicing intron (2R8S); **(G)** T-loop/purine interaction from yeast tRNA^phe^ involving three chemically modified nucleotides (1EHZ); **(H)** RNA quadruplex including an inosine tetrad (2GRB). Colors indicate accurately recovered noncanonical features (pastel colors), accurately recovered extrahelical bulges (wheat with white side-chains), flanking helices built *de novo* (violet), parts of experimental structure used for modeling but allowed to minimize (dark violet), fixed context from experimental structure (black in 2D; white in 3D), additional helical context not included in modeling (gray in 2D; white in 3D).

**Table 1.**
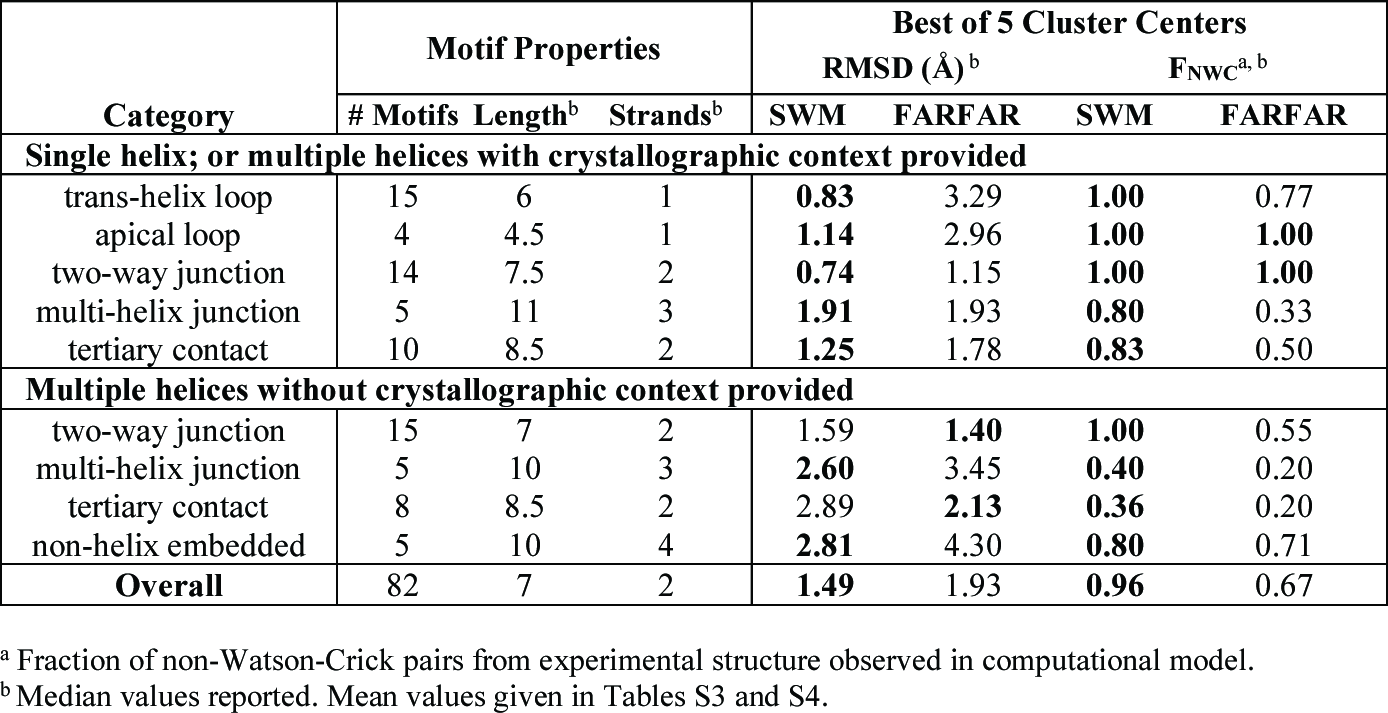
Benchmark of stepwise Monte Carlo (SWM) compared to prior Rosetta fragment assembly method (FARFAR) over different classes of RNA structure motifs.

Perhaps the most striking models were recovered for multi-helix junctions and tertiary contacts, which have largely eluded RNA modeling efforts seeking high resolution (*6, 8*). SWM achieves high accuracy models for the P2-P3-P6 three-way junction from the Varkud satellite ribozyme, previously missed by all modelers in the RNA-puzzle 7 challenge (Fig. 2D; 1.13 Å RMSD, 3 of 3 noncanonical pairs recovered); a highly irregular tertiary contact in a hammerhead ribozyme (Fig. 2E; 1.16 Å RMSD, 2 of 3 noncanonical pairs and 1 extrahelical bulge recovered); a complex between a GAAA tetraloop and its 11-nt receptor (0.64 Å RMSD, all 4 noncanonical pairs recovered when flanking helix context was provided, Fig. 2F); and the tRNA^phe^ T-loop, a loop-loop tertiary contact stabilized by chemical modifications at 5-methyl-uridine, pseudouridine, and N1-methyl adenosine (Fig. 2G; 1.33 Å accuracy when flanking context was provided). Motifs without any flanking A-form helices offered particularly stringent tests for *ab initio* modeling and could also be recovered at high accuracy by SWM, as illustrated by the inosine-tetrad-containing quadruplex (Fig. 2H; 2.87 Å RMSD overall, 0.46 Å RMSD if the terminal uracils, which make crystal contacts, are excluded). For comparison, we also carried out modeling with FARFAR on these 82 motifs, taking care to remove possible homologs from the method’s fragment library to mimic a realistic *ab initio* prediction scenario (Supplementary Methods; we could not carry out fair comparisons to other methods due to the unavailability of similar homolog exclusion options in those methods). SWM strongly outperformed these FARFAR models in terms of recovery of noncanonical pairs (*p* < 5 × 10^−5^) and RMSD accuracy (*p* < 2 × 10^−4^) (Figure S3, Table 1, Table S3, and Table S4; *p* values are based on Wilcoxon ranked-pairs test, *n* = 82).

In some benchmark cases, SWM did not exhibit near-atomic-accuracy recovery and illuminated challenges remaining for computational RNA modeling. While a few discrepancies between SWM models and X-ray structures could be explained by crystallographic interactions (e.g., edge nucleotides making crystal contacts, Fig. 3H), the majority of problems were better explained by errors in the energy function. For 9 of the 14 cases in which the SWM modeling RMSD was worse than 3.0 Å (and thus definitively not achieving atomic accuracy), the energy of the lowest energy SWM model was lower than that of the optimized experimental structure, often by several units of free energy (calibrated here to correspond to k_B_T (*20*); Table S3). One hint for the issues in the energy function came from cases where the fragment-based method (FARFAR) outperformed SWM if assessed by RMSD but not if assessed the fraction of base pairs recovered (Table 1). The existence of such FARFAR models with native-like backbones but incorrect base pairs suggested that conformational preferences implicitly encoded in database fragments in FARFAR might needed to be better captured during SWM. One possible route to improving SWM might be to update the RNA torsional potential of the Rosetta free energy function, which currently does not model correlations across most backbone torsions. Results on the hepatitis C virus internal ribosome entry site, the sarcin ricin loop, and other test cases suggest that a modified torsional potential, as well as inclusion of metal ions, may eventually address these residual problems (Fig. S4).

**Figure 3.**
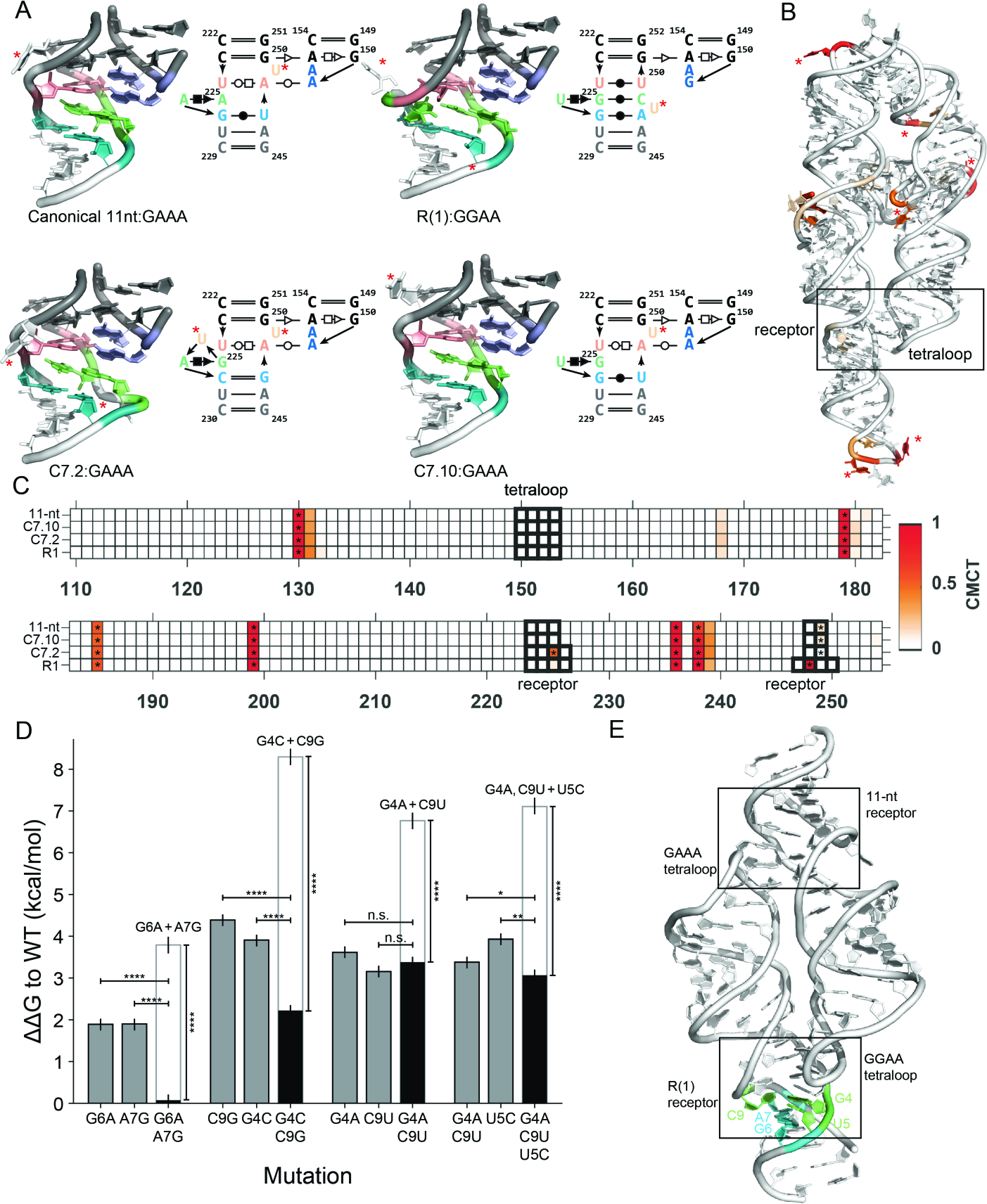
SWM modeling and prospective experimental tests of previously unsolved tetralooptetraloop receptor (TL/TR) motifs. **(A)** *Ab initio* SWM models for canonical 11-nucleotide TL/TR motif and alternative motifs discovered through *in vitro* selection that have resisted crystallization. Lavender, salmon, lime, and teal colorings highlight homologous structural features; during modeling, bottom flanking helix (white) was allowed to move relative to the top helices of the receptor and tetraloop (gray), which were held fixed. **(B)** The canonical 11-nt tetraloop receptor module from the P4-P6 domain of the *Tetrahymena* group I self-splicing intron (PDB: 2R8S). In **(A-B)**, red asterisks mark uracil residues predicted to be bulged. (C) CMCT mapping of the receptors installed into the P4-P6 domain of the *Tetrahymena* ribozyme (tetraloop and receptor indicated by black boxes) supports the bulged uracils in the predicted models (black asterisks). **(D)** Selective tests of each R(1) receptor base pair by compensatory mutagenesis in tectoRNA dimer. Rescue by double and triple mutants (black bars) was compared to energetic perturbations predicted based on sum of effects (white bars) of component mutations or, more conservatively, to the single mutants. One, two, three, and four asterisks represent *p* values (computed by Student’s t-test for difference of means) less than 0.05, 0.001, 5 × 10^−5^, and 1 × 10^−6^, respectively; n.s., not significant. **(E)** Overall 3D model of tectoRNA dimer with SWM model for R(1) receptor.

### Stringent tests of SWM models for new RNA-RNA tertiary contact motifs

As we began to see significant improvements of modeling accuracy in the 82-motif benchmark, we hypothesized that SWM might be able to predict noncanonical base pairs in motifs that have been refractory to NMR and crystallographic analysis. Success in the 11-nt tetraloop/receptor benchmark test case (Fig. 2F), a classic model system and ubiquitous tertiary contact in natural RNAs, encouraged us to model alternative tetraloop/receptor complexes selected for use in RNA engineering but not yet solved experimentally (*2, 22*), and to design stringent experimental tests to validate or falsify these models.

A detailed sequence/function analysis previously suggested similarities between the GAAA/C7.2, GAAA/C7.10, and GGAA/R(1) interactions discovered through *in vitro* evolution (*22*) and the classic GAAA/11-nt receptor, which has been crystallized in numerous contexts. It has not been clear however if this similarity holds at the structural level due to the unavailability of high resolution structures of the three artificial tetraloop/receptors. For example, prior literature analyses conflicted in the proposal of which C7.2 receptor nucleotides, if any, might form a ‘platform’ (lime, Fig. 3A), analogous to A4-A5 platform in the 11-nt receptor (G4 and A6 with an intervening bulge in (*11, 23*); and G4 and U5 in (*22*)). Similarly, a proposed homology of C9 in R(1) to A8 in the 11-nt receptor (*22*) has not been tested by structure modeling or experiments.

We carried out stepwise Monte Carlo modeling to explore possible structural homologies of these four receptors. In the SWM runs, the stem and basal G-A sugar-Hoogsteen pair of the GNRA tetraloop and their docking site into the GG/CC stem of the receptor were seeded, based on one crystal structure of the GAAA/11-nt receptor (see considerations of A-minor geometries discussed in (*22*)). The remaining 10 nucleotides and receptor stem were modeled *ab initio*. SWM modeling of all four of these receptors achieved convergence, with 8 of the top 10 models clustered within 2 Å RMSD of each other. The modeling recovered the known GAAA/11-nt structure at 0.80 Å RMSD and reproduced a previous C7.2 model that involved SWA enumeration of only three nucleotides (G4-U5-A6) rather than rebuilding *ab initio* the complete tetraloop/receptor interaction.

SWM models for all four tetraloop/receptors exhibited striking structural homology to each other but also noncanonical features (extrahelical bulges, pairs) that were not anticipated from prior manual modeling efforts (Fig. 3A; Supplemental Text; models provided in Supplemental File). Three features were preserved across loops. First, models for all receptors exhibited a docking site for the second nucleotide of the tetraloop (salmon, Fig. 3A). In GAAA/11-nt, GAAA/C7.2, and GAAA/C7.10, where the second tetraloop nucleotide is A, the receptor docking site was predicted to be an adenosine that is part of a Watson-Crick/Hoogsteen U-A pair. In GGAA/R(1), where the second tetraloop nucleotide is G, the receptor docking site was predicted to be U3, part of a noncanonical Watson-Crick U3-U10 pair. Second, SWM models for all receptors exhibited a platform involving two same-strand base-paired nucleotides that stack under the tetraloop (lime, Fig. 3A). The sequence varies, however, between A-A in the 11-nt receptor; G-U in the C7.10 receptor; G-A in the C7.2 receptor (supporting the models in (*11, 23*) but not the model in (*22*)); and G-U in the R(1) receptor. In the R(1) receptor, C9 was predicted by SWM to form a stabilizing C-G base pair with a platform nucleotide. While C9 was previously proposed to be homologous to A8 in the 11-nt receptor (*22*), the new model also explains prior binding data that implicated C9 as forming core interactions in R(1). Third, all receptors show a noncanonical pair involving Watson-Crick edges, needed to transition between the platform region and the lower stem of the receptor (teal, Fig. 3A). The sequence is a G-A pair in R(1), and a G•U wobble in the others. Overall, given the sequence mapping between receptors revealed by the SWM models, each noncanonical pairing in the naturally occurring GAAA/11-nt structure had a homolog, albeit one that was difficult to predict (and, in some cases, differently predicted) in each of the three non-native tetraloop receptors.

We tested these features using prospective experiments. The SWM models predicted different single uridines to be bulged out of each tetraloop receptor. Reaction to CMCT (*N*-cyclohexyl-*N*′-(2-morpholinoethyl) carbodiimide tosylate) followed by reverse transcription allows single-nucleotide-resolution mapping of unpaired uridines that bulge out of structure and expose their Watson-Crick edges to solution. We therefore installed the tetraloop/receptors into the P4-P6 domain of the *Tetrahymena* ribozyme (Fig. 3B), which also displays other bulged uridines that served as positive controls (asterisks in Fig. 3C). These experiments verified extrahelical bulging of single-nucleotide uridines predicted by SWM at different positions in the different receptors, and disfavored prior manual models (Fig. 3C and Supplemental Text).

We carried out further prospective experiments to incisively test base pairs newly predicted by SWM modeling. In particular, the R(1) receptor model included numerous unexpected noncanonical features, especially a base triple involving a new Watson-Crick singlet base pair G4-C9 and a dinucleotide platform at G4-U5. These features were stringently evaluated via compensatory mutagenesis. Chemical mapping on the P4-P6 domain confirmed the G4-C9 base pair but was not sensitive enough to test other compensatory mutants (Fig. S5). We therefore carried out native gel assembly measurements in a different system, the tectoRNA dimer, which enables precise energetic measurements spanning 5 kcal/mol (Figs. 3D-E). Observation of energetic disruption by individual mutations and then rescue by compensatory mutants confirmed the predicted interactions of G4-C9, the base triple G4-U5-C9, and noncanonical pair G6-A7 (*p* < 0.01 in all cases; Fig. 3D) as well as other features of the model (Supplemental Text and Fig. S6). Overall, these experimental results falsified bulge predictions and base pairings previously guessed for these tetraloop/receptors (*11, 22, 23*) and strongly supported the models predicted by SWM. Our structural inference and mutagenesis-based validation of noncanonical pairs would have been intractable without the SWM-predicted models, due to the large number of possible mutant pair and triple combinations that would have to be tested.

### Blind prediction of all non-canonical pairs of a community-wide RNA-puzzle

The community-wide modeling challenge RNA-Puzzle 18 provided an opportunity to further blindly test SWM and to compare it to best efforts of other state-of-the-art algorithms (Fig. 4). This problem was of mixed difficulty. On one hand, the 71-nucleotide target sequence was readily identified via PDB-BLAST (*24*) to be a Zika virus RNA homologous to a molecule with a previously solved X-ray structure, an Xrn1 endonuclease-resistant (xrRNA) fragment of Murray Valley Encephalitis virus (PDB ID: 4PQV) (*25*). However, the crystallographic environment of the prior structure disrupted a pseudoknot (between L3 and J1/4, Fig. 4A) expected from sequence alignments so that nearly half of the prior structure could not be trusted as a template for homology modeling. Intermolecular crystal contacts produced an open single-stranded region in the asymmetric unit where the pseudoknot was expected and interleaved regions from separate molecules; the scale of these conformational perturbations was as large as the dimensions of the molecule itself (Fig. S7). Further complicating the modeling, two Watson-Crick pairs within stem P3 changed to or from G•U wobble pairs. Moreover, prior literature analysis (*25*) suggested extension of this helix by two further Watson-Crick pairs (U29-A37; U30-A36), albeit without direct evidence from phylogenetic covariation and in partial conflict with dimethyl sulfate probing. *Ab initio* modeling at a scale inaccessible to the prior enumerative SWA method was necessary for modeling the RNA, and we therefore carried out stepwise Monte Carlo (Fig. 4B; Methods).

**Figure 4.**
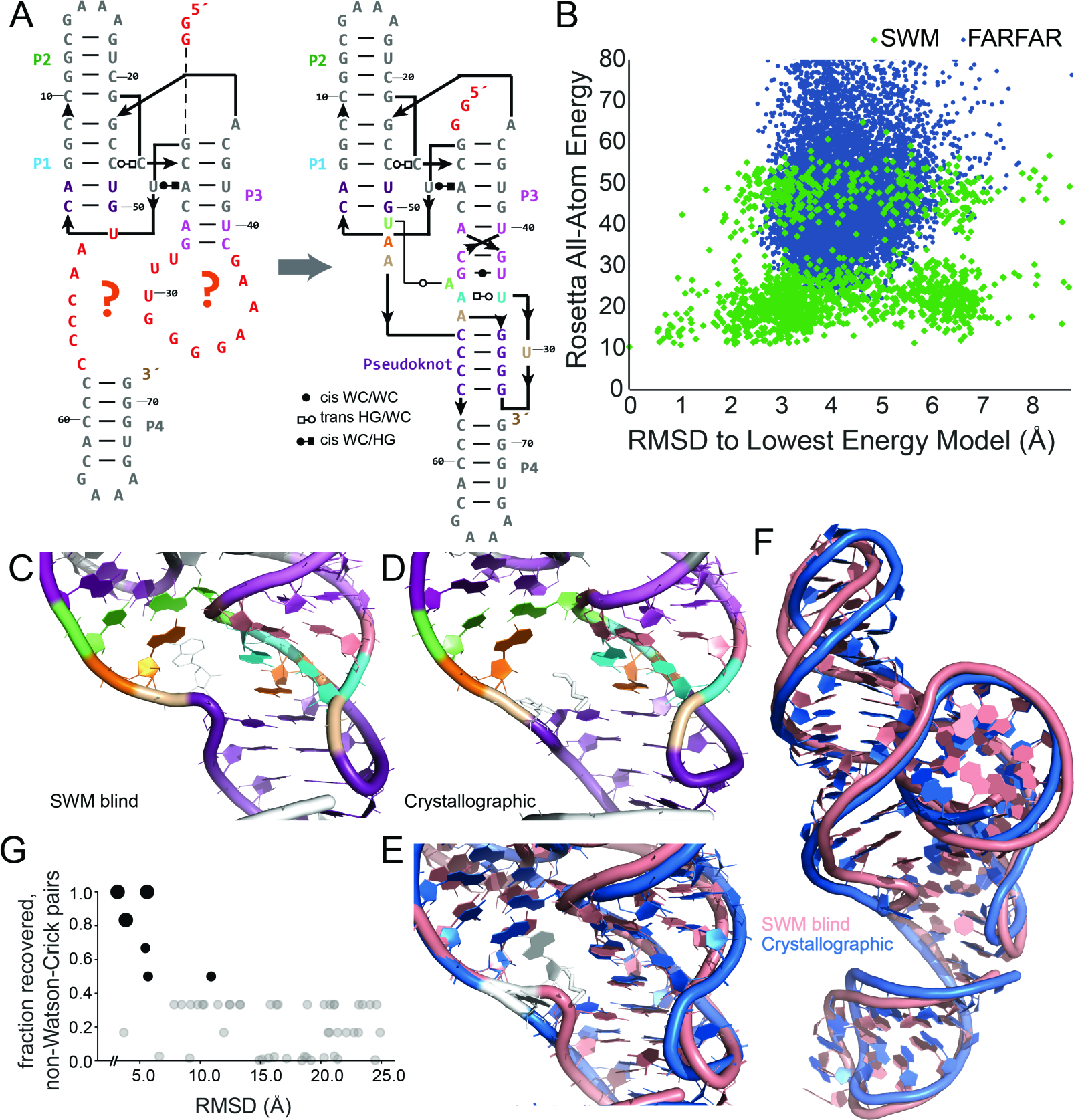
Blind prediction of complex RNA tertiary fold during RNA-Puzzle 18. **(A)** Two-dimensional diagram of the RNA Puzzle 18 (Zika xrRNA) modeling problem, highlighting motifs that needed to be build *de novo* in red (left) and SWM-predicted pairings (pastel colors, right). **(B)** Structures discovered by stepwise Monte Carlo (SWM, green) are lower in energy and ˜ 4 Å from models in conventional fragment assembly (FARFAR, blue); note x-axis is RMSD to lowest free energy SWM model, not the experimental structure (unavailable at time of modeling). Magnified view of noncanonical region built *de novo* for **(C)** SWM model submitted for RNA-Puzzles competition and **(D)** the subsequently released crystal structure; **(E)** and **(F)** give overlays in magnified and global views, respectively (SWM, salmon; crystal, marine). **(G)** Fraction of noncanonical base pairs recovered and RMSD to native model obtained by Rosetta modeling (black; larger and smaller symbols are SWM and FARFAR, respectively) and other labs (gray) for RNA-Puzzle 18. Points recovering zero noncanonical pairs are given a small vertical perturbation to appear visually distinct.

The lowest free energy SWM models for RNA-puzzle 18 converged to a tight ensemble of intricate structures, with one submitted SWM model shown in Fig. 4C. The Watson-Crick pairs U29-A37 and U30-A36 predicted in the literature did not occur in the models. Instead, several other features were consistently observed across the SWM models (colored in Fig. 4A, right panel and Fig. 4C): co-axial arrangement of the pseudoknot helix (purple) on P3 (light violet), a noncanonical *trans* Watson-Crick base pair between A37 and U51 stacking under P1 (green), a UA-handle (*26*) formed by U29-A36 (turquoise), and lack of pairing by U30, A35, A52, and A53 (sand and orange). These features were not uniformly present – or not predicted at all – in models created by FARFAR or, as it later turned out, in models submitted by other RNA-Puzzle participants (Fig. S8).

The subsequent release of the crystal structure (Figs. 4D and E) (*27*) confirmed all base pairs predicted by SWM modeling (100% non-Watson-Crick recovery). Indeed, the only structural deviation involved A53 (sand, Fig. 4C), which was correctly predicted in SWM models to be unpaired but also stacked on neighbor A52 (orange, Fig. 4C). In the crystal, A53 was indeed unpaired but bulged out of the core to form a contact with a crystallographic neighbor, while a 1,6-hexanediol molecule from the crystallization buffer took its place (white sticks, Fig. 4C); this arrangement was noted independently to be a likely crystallographic artifact (*27*). There is striking overall fold agreement (3.08 Å RMSD; and 1.90 Å over just the most difficult noncanonical region, nucleotides 5-6, 26-40, 49-59, and 70-71; Figs. 4C and 4D), much better than the ˜10 Å best-case agreement seen in previous RNA-puzzles of comparable difficulty (*2*). Furthermore, SWM predicted all noncanonical base pairs accurately (F_NWC_ = 1, Fig. 4G). While one blind model from another method achieved somewhat comparable RMSD to the crystal structure (3.61 Å), it predicted only one of six non-Watson-Crick base pairs (Fig. 4G) and left a ‘hole’ in the central noncanonical region (RMSD of 3.67 Å in that region, Fig. S8).

## Discussion

We have presented an algorithm for modeling RNA structures called stepwise Monte Carlo, which uniquely allows for the addition and deletion of residues during modeling guided by the Rosetta all-atom free energy function. The minima of the energy landscape are efficiently traversed by this method, allowing the *ab initio* recovery of small RNA loop structures in hours of CPU-time (Fig. 1). On an extensive benchmark, SWM enables quantitative recovery of noncanonical pairs in cases that include prior RNA-puzzle motifs, junctions and tertiary contacts involving numerous strands, and motifs without any A-form helices (Fig. 2, Table 1). We applied SWM to model structures of three previously unsolved tetraloop-receptors and prospectively validated these models through chemical mapping and extensive compensatory mutagenesis (Fig. 3). Last, SWM achieved blind prediction of all noncanonical pairs of a recent RNA-puzzle, an intricately folded domain of the Zika RNA genome whose pairings were missed by other methods applied by our and other modeling groups (Fig. 4). The most striking aspect of the SWM models is the high recovery of noncanonical pairs, which have largely eluded prior algorithms when tested in blind challenges. These results support stepwise nucleotide structure formation as a predictive algorithmic principle for high resolution RNA structure modeling, and we expect SWM to be useful in the *ab initio* modeling and refinement of noncanonical RNA structure.

The results above focused on solving individual noncanonical motifs. While such problems arise frequently in real-world modeling (e.g., the unsolved tetraloop receptors), most functional RNA structures harbor multiple junctions and tertiary contacts whose folds become dependent on each other through the lever-arm like effects of interconnecting helices. SWM is currently too computationally expensive to simultaneously simulate all motifs and helices in such molecules. It may be necessary to better parallelize the current algorithm to allow concomitant modeling of multiple motifs on multi-processor computers, as is routine in molecular dynamics simulations (*28*). Alternatively, modeling may benefit from iterating back-and-forth between high-resolution SWM and complementary low-resolution modeling approaches like MC-Sym/MC-Fold, Rosetta FARFAR, iFoldRNA, SimRNA, and VFold3D (*6-10*), similar to iterative approaches in modeling large proteins (*5*). In addition, we note that SWM relies heavily on the assumed free energy function for folding, and several of our benchmark cases indicate that even the most recently updated Rosetta free energy function is still not accurate when SWM enables deep sampling. Therefore, a critical open question is whether residual free energy function problems might be corrected by improved RNA torsional potentials, treatment of electrostatic effects, or by use of energy functions independently developed for biomolecular mechanics and refinement (*5, 28, 29*).

## Materials and Methods

### Stepwise Monte Carlo

Stepwise Monte Carlo (SWM) was implemented in C++ in the Rosetta codebase. The source code and the *stepwise* executable compiled for different environments are being made available in Rosetta release 3.6 and later releases, free to academic users at http://www.rosettacommons.org. Full documentation, including example command-lines, tutorial videos, and demonstration code, is available at https://www.rosettacommons.org/docs/latest/application_documentation/stepwise/stepwise_monte_carlo/stepwise.

The full set of benchmark cases, including the 82 central to this work, are available at https://github.com/DasLab/rna_benchmark. The repository contains input files for each benchmark case; scripts for setting up benchmark runs using either stepwise Monte Carlo or fragment assembly, including automated job submission for multiple cluster job schedulers; and scripts for creating analysis figures and tables. Finally, SWM is available through a web server on ROSIE, the Rosetta Online Server that Includes Everyone, at https://rosie.rosettacommons.org/stepwise.

**Supplementary Methods** gives detailed descriptions of SWM, stepwise assembly (SWA), and fragment assembly of RNA with full-atom refinement (FARFAR) modeling; evaluation of RMSD and energetic sampling efficiency; and Protein Databank accession IDs for experimental structures.

### Chemical mapping

Chemical mapping was carried out as in (*30*). Briefly, DNA templates for the P4-P6 RNA were produced through PCR assembly of oligonucleotides of length 60 nucleotides or smaller (Integrated DNA Technologies) using Phusion polymerase (Finnzymes, MA). DNA templates were designed with the T7 RNA polymerase promoter (TTCTAATACGACTCACTATA) at their 5´ ends. A custom reverse transcription primer-binding site (AAAGAAACAACAACAACAAC) was included at the 3´ terminus of each template. RNA transcribed with T7 RNA polymerase (New England Biolabs) was purified using RNAClean XP beads (Beckman Coulter). RNA modification reactions were performed in 20 µL reactions containing 1.2 pmol RNA. RNAs were incubated with 50 mM Na-HEPES (pH 8.0) at 90 °C for 3 min then cooled to room temperature. MgCl_2_ at 0 or 10 mM final concentration was then added, followed by incubation at 50 °C for 30 minutes and then room temperature prior to chemical mapping. Chemical probes were used at the following final concentrations: DMS (0.125% v/v), CMCT in water (2.6 mg/mL), 1M7 (1.05 mg/mL in anhydrous DMSO gave final DMSO concentration of 25%). Chemical probes were allowed to react for 15 minutes prior to quenching. 1M7 and CMCT reactions were quenched with 5.0 µL of 0.5 M Na-MES (pH 6.0), while the DMS reaction was quenched with 3.0 µL of 3 M NaCl, 1.5 µL of oligo-dT beads (poly(A) purist, Ambion), and 0.25 µL of a 0.25 µM 5´-FAM-A20-Tail2 primer which complements the reverse transcription primer-binding site at the RNA 3´ ends. The quench mixture was incubated at room temperature for 15 min and the purification beads were pulled down with 96 post magnetic stand, washed with 100 µL of 70% ethanol twice for RNA purification. RNAs were reverse transcribed with Superscript III Reverse Transcriptase at 48 °C for 40 min (Life Technologies). RNA template was subsequently hydrolyzed for 3 min at 90 °C in 0.2 M NaOH. After pH neutralization, cDNA on Oligo dT beads was pulled down by magnetic stand and perform ethanol wash as above. cDNAs were eluted into 10 µL of Rox350 standard ladder in HiDi-formamide (Life Technologies), using 1 µL of Rox350 in 250 µL of HiDi-formamide. ABI3700 sequencers were used for electrophoresis of cDNA. CE data were quantitated with HiTRACE (*31*). Data from these P4-P6 RNA experiments have been posted to the RNA Mapping Database (*32*) at the following accession IDs:

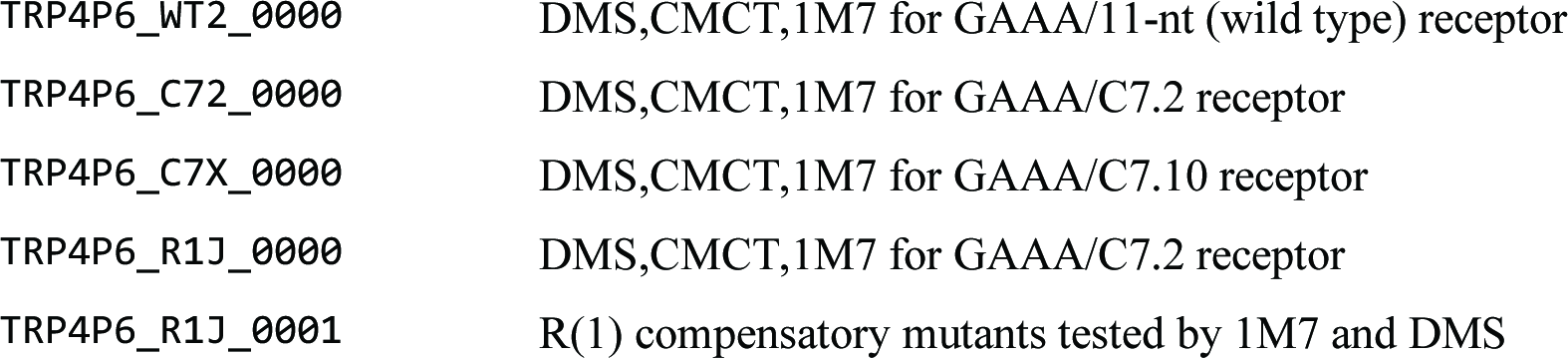

### Native gel shift experiments

Gel shifts were performed as previously described (*22*). Briefly, equimolar amounts of each RNA monomer at various concentrations (up to 20 µM final) were mixed in water and denatured at 95°C for 1 min. Mixtures were cooled on ice for 2 min and annealed at 30 °C for 5 min before addition of Mg^2+^ buffer [(9 mM Tris–borate pH 8.3, 15 mM Mg(OAc)_2_ final concentration). After 30 min incubation at 30 °C, samples were incubated at 10 °C for 15 min before native gel analysis (7% (29:1) polyacrylamide gels in Mg^2+^ buffer at 10°C). One of the monomers contained a fixed amount of 3′ end [^32^P] pCp-labeled RNA (∼0.25–0.5 nM final). Monomer and dimer bands were quantified with ImageQuant and dimer formation was plotted against RNA concentration. *K*_d_’s were determined as the concentration at which half the RNA molecules dimerized, and converted to DG (relative to 1 M standard state) through the formula DG = *k*_B_T ln(*K*_d_ / 1 M), where k_B_T is the Boltzmann constant and *T* is the temperature.

## General

We thank F.-C. Chou for early discussions; and Stanford Research Computing for expert administration of the BioX^3^ clusters (supported by NIH 1S10RR02664701) and Sherlock clusters.

## Author contributions

A.W., C.G., L.J., and R.D. designed research; A.W., C.G., W.K., P.Z., R.D. carried out research; A.W., L.J., R.D. wrote the manuscript; all authors reviewed the manuscript.

**Data and materials availability:** all data and computer code needed to evaluate the conclusions in this paper are available at links described in Materials and Methods.

